## Supplementary Figure 1 for "A large-scale cancer-specific protein-DNA interaction network"

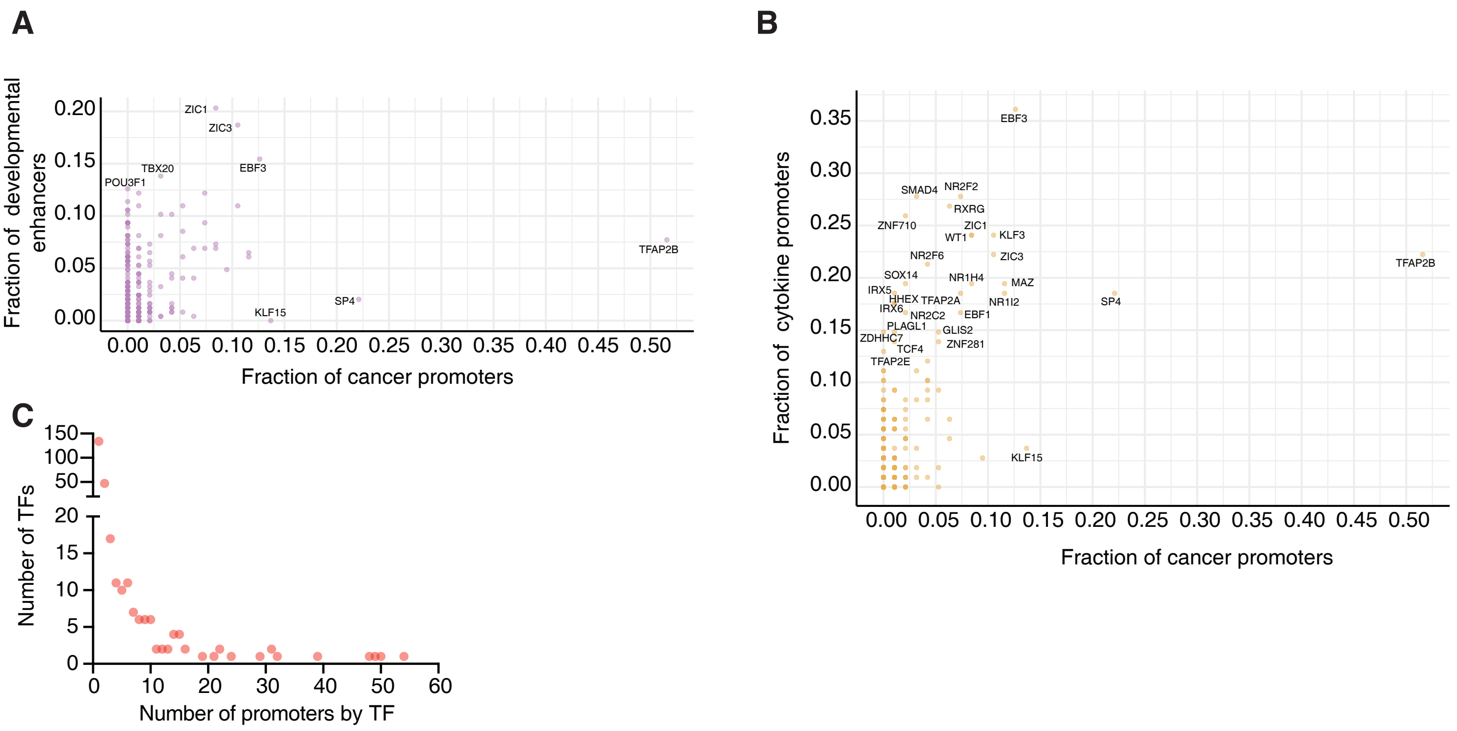


**Supplementary Figure 1: Comparison between the TF connectivity in different networks.** (**A-B**) Percentage of protein-DNA interactions for each TF in the cancer gene network versus the developmental enhancer (**A**) or the cytokine gene (**B**) networks. (**C**) Distribution of the number of TFs with the binding the indicated number of gene promoters.
